# Selective sweeps identification in distinct groups of cultivated rye (*Secale cereale* L.) germplasm provides potential candidates for crop improvement

**DOI:** 10.1101/2023.01.22.525081

**Authors:** Anna Hawliczek, Ewa Borzęcka, Katarzyna Tofil, Nikolaos Alachiotis, Leszek Bolibok, Piotr Gawroński, Dörthe Siekmann, Bernd Hackauf, Roman Dušinský, Miroslav Švec, Hanna Bolibok-Brągoszewska

**Affiliations:** Department of Plant Genetics Breeding and Biotechnology, Institute of Biology, Warsaw University of Life Sciences-SGGW, Warsaw, Poland; Faculty of Electrical Engineering, Mathematics and Computer Science, University of Twente, Enschede, The Netherlands; Department of Silviculture, Institute of Forest Sciences, Warsaw University of Life Sciences-SGGW, Warsaw, Poland; HYBRO Saatzucht GmbH and Co. KG, Schenkenberg, Germany; Julius Kühn-Institut, Groß Lüsewitz, Germany; Department of Botany, Faculty of Natural Sciences, Comenius University in Bratislava, Bratislava, Slovakia

**Keywords:** rye, *Secale cereale* L, selective sweeps, genetic diversity, population structure, GBS, DArTseq

## Abstract

**Background:** During domestication and subsequent improvement plants were subjected to intensive positive selection for desirable traits. Identification of selection targets is important with respect to the future targeted broadening of diversity in breeding programmes. Rye (*Secale cereale* L.) is a cereal that is closely related to wheat, and it is an important crop in Central, Eastern and Northern Europe. The aim of the study was (i) to identify diverse groups of rye accessions based on high-density, genome-wide analysis of genetic diversity within a set of 478 rye accessions, covering a full spectrum of diversity within the genus, from wild accession to inbred lines used in hybrid breeding, and (ii) to identify selective sweeps in the established groups of cultivated rye germplasm and putative candidate genes targeted by selection.

**Results:** Population structure and genetic diversity analyses based on high-quality SNP (DArTseq) markers revealed the presence of three complexes in the *Secale* genus: *S. sylvestre, S. strictum* and *S. cereale*/*vavilovii*, a relatively narrow diversity of *S. sylvestre*, very high diversity of *S. strictum*, and signatures of strong positive selection in *S. vavilovii*. Within cultivated ryes we detected the presence of genetic clusters and the influence of improvement status on the clustering. Rye landraces represent a reservoir of variation for breeding, and especially a distinct group of landraces from Turkey should be of special interest as a source of untapped variation. Selective sweep detection in cultivated accessions identified 133 outlier positions within 13 sweep regions and 170 putative candidate genes related, among others, to response to various environmental stimuli (such as pathogens, drought, cold), plant fertility and reproduction (pollen sperm cell differentiation, pollen maturation, pollen tube growth),and plant growth and biomass production.

**Conclusions:** Our study provides valuable information for efficient management of rye germplasm collections, which can help to ensure proper safeguarding of their genetic potential and provides numerous novel candidate genes targeted by selection in cultivated rye for further functional characterisation and allelic diversity studies.

## BACKGROUND

During domestication and subsequent diversification and improvement plants were subjected to intensive positive selection. Consequently, several key traits differentiate crop plant from their wild progenitors. In the case of cereal crops, these traits include: larger grain size, loss of natural seed dispersal mechanisms (causing seed retention until harvest), changes in THE plant’s architecture (apical dominance), and in plant physiology (changes related to seed dormancy, photoperiodic sensitivity, vernalization requirements) [1, 2]. A number of genes responsible for domestication traits had been already identified and characterized in major crops, such as maize, rice or wheat, for example *Q* (controlling inflorescence structure in wheat), *teosinte branched1* (*tb1*, controlling shoot architecture in maize), *Shattering1* (*Sh1*, causing the loss of seed shattering), *Btr1* and *Btr2* (required for the disarticulation of rachis)[1, 3, 4]. Diversification genes, targeted by selection after domestication, are responsible for intervarietal differences and are typically related to yield, biotic and abiotic stress resistance, grain quality and adaptation [2]. Well know examples of such genes are: maize *Y1* gene, related to high carotenoids levels and yellow kernels [5], wheat *Rht* gene, controlling reduced height, and rice *Hd1*, controlling flowering time [5–7].

At first, the QTL approach was predominately used to identify domestication/improvement loci. More recently, various population genetic approaches were developed to detect selective sweeps based on genome-wide scans, including population differentiation and environmental association methods [7–11].

In contrast to early studies, suggesting that several, large effect loci underlie the phenotypic switch from a wild progenitor to a domesticate, genome-wide studies revealed hundreds of loci showing signatures of selection [8, 12, 13], providing a new insight into the influence of domestication and breeding on the genome and numerous potential candidate genes for crop improvement programs. Nevertheless, despite extensive research on the subject, the knowledge of the influence of domestication and improvement on the genome is still very incomplete in many crops.

Rye (*Secale cereale* L.) is a cereal closely related to wheat, and an important crop in Central, Eastern and Northern Europe. It is mainly used for the production of flour for bread making, as animal feed, and in distilleries to produce whiskey and vodka. Rye has the highest tolerance of abiotic and biotic stresses (including cold temperature, low soil fertility, and high soil acidity) among the small grain temperate cereals and is a widely used source of genetic variation for wheat improvement [14, 15]. Contrary to its closest crop relatives wheat and barley, cultivated rye is outcrossing. Recently genome sequences of two rye accessions were published [16, 17]. Rye genome size ranges from 7.68 to 8.03 Gbp, and the repetitive elements account for 85%-90% of the assemblies.

According to Germplasm Resource Information Network (GRIN) there are four species recognized in the *Secale* genus: *S cereale, S strictum, S vavilovii, S sylvestre*. Many molecular studies indicate, however, that S. *vavilovii* is a part of *S cereale* complex, and postulate a revision of *Secale* classification [18–21]. A possible explanation for the discrepancies regarding classification of *S. vavilovii* was provided by Zohary et al. [22], who proposed that the four complexes within *Secale* are *cereale, strictum, iranicum*, and *sylvestre*. “True*” S. vavilovii* forms belong to *S. cereale* complex, which is supported be extensive molecular data mentioned above. *S. iranicum* (Kolbylansky) is poorly know, and was at a point of time erroneously described as *vavilov*ii and sent to several germplasm collection under this description causing confusion, with some researches working on the ‘true’ *vavilovii*, and others on *iranicum*, only mistakenly described as *vavilovii*. Thus, the matter of *Secale* classification is not fully resolved yet.

Rye domestication happened approximately four thousand years ago [22, 23], much later than the domestication of wheat or barley (ca. 10 thousand years ago [24]. Prior to that, rye occurred as a weed in wheat and barley cultures [25]. For these reasons, rye is referred to as a secondary domesticate [23]. There is no consensus regarding the immediate wild progenitor of cultivated rye (*S. cereale* subsp. *cereale*), with *S. vavilovii* and *S. strictum*, among others, being suggested as likely candidates [23, 26]. Central and Eastern Turkey and adjacent regions are reported to be the main centre of diversity of rye wild species [25]. Recent genetic diversity scans indicate, that there is considerable diversity within rye genetic resources and that the current breeding pool is genetically relatively narrow and distant from accessions representing genebank collections. Additionally, no clear correspondence of genetic diversity patterns with geographic origins was observed [18, 27–30].

The aim of this study was to: i) assess the genetic diversity structure in a diverse collection of 478 rye accessions representing different geographic origins and improvement status based on genome-wide, high-quality GBS markers, ii) identify selective sweeps in established germplasm clusters and iii) indicate potential candidate genes targeted by selection in rye.

## RESULTS

### GBS (DArTseq) genotyping

In total 79 877 SNP markers (Dataset-1) differentiating 478 rye accessions (Table S1) were identified and 49 977 (62.65%) of them could be aligned to the Lo7 rye reference genome sequence [16], with the markers fairly evenly distributed among chromosomes - the percentage of SNPs mapped to individual chromosomes ranged from 11.3 for 1R to 15.6 for 2R. After quality filtering, 12 846 high quality (HQ) SNP markers (Dataset-2) were identified and used for population structure and phylogenetic analyses (Table S2). Of those, 10 607 (82.6%) aligned to the Lo7 genome sequence and spanned 99.7 % of the assembly (6.72 Gb). Percentage of HQ SNP markers mapped to individual chromosomes varied between 11.6 for 1R to 17.4 for 2R (Table S3). The average MAF and PIC values for both all 12 846 HQ SNPs and 10607 HQ SNPs mapped to genome sequence were 0.15 and 0.17, respectively. Distribution of MAF and PIC values of 12846 HQ SNPs is shown in Figure S1.

### Population structure

#### Assignment tests

K=2 was found to explain best the population structure (Figure S2). Using the cut-off value of Q ≥ 70 %, 430 and 34 accessions were assigned to populations 1 and 2, respectively, while 13 accessions were classified as admixtures (Figure S3, Table S1). Population 1 comprised all accessions representing *S. vavilovii* and *Secale cereale* ssp. included in the study (only one *S. c*. subsp. *dighoricum* accession was assigned to population 2), and 14 accessions described as unknown in genebank records. Five *S. s*. subsp. *strictum* and three *S. s*. subsp. *anatolicum* accessions were also assigned to population 1 - ca. 24% and 38% of accessions representing these taxa included in the study, respectively. Population 2 included all ten *S. sylvestre* accessions and accessions representing *S. strictum* subspecies: most of *S. s*. ssp. *kuprijanovii* accessions (11 accessions, 79 %) and also accessions of *S. s*. ssp. *strictum* (nine accessions, 43%) and *S. s*. ssp. *anatolicum* (three accessions, 38%). The remaining three *S. s*. ssp. *kuprijanovii* accessions were classified as admixtures, together with seven *S. s*. ssp. *strictum*, two *S. s*. subsp. *anatolicum*, and two *S. s*. subsp. *africanum* accessions.

### Principal Coordinates Analysis

PCoA and STRUCTURE results were in a very good agreement, however, PCoA revealed a more complex structure in the germplasm set by clustering the accessions into three groups (Figure 1). Accessions assigned to population 2 by STRUCTURE were divided into two groups in the PCoA plot – a group containing *S. sylvestre* accessions and a group containing *S. strictum* accessions. The third group indicated by PCoA, occupying a relatively small area of diversity space, corresponded to population 1 indicated by STRUCTURE. As expected, landraces were dispersed across a larger plot area than modern and historical cultivars, and thus turned out to be more diverse.

**Figure 1.**
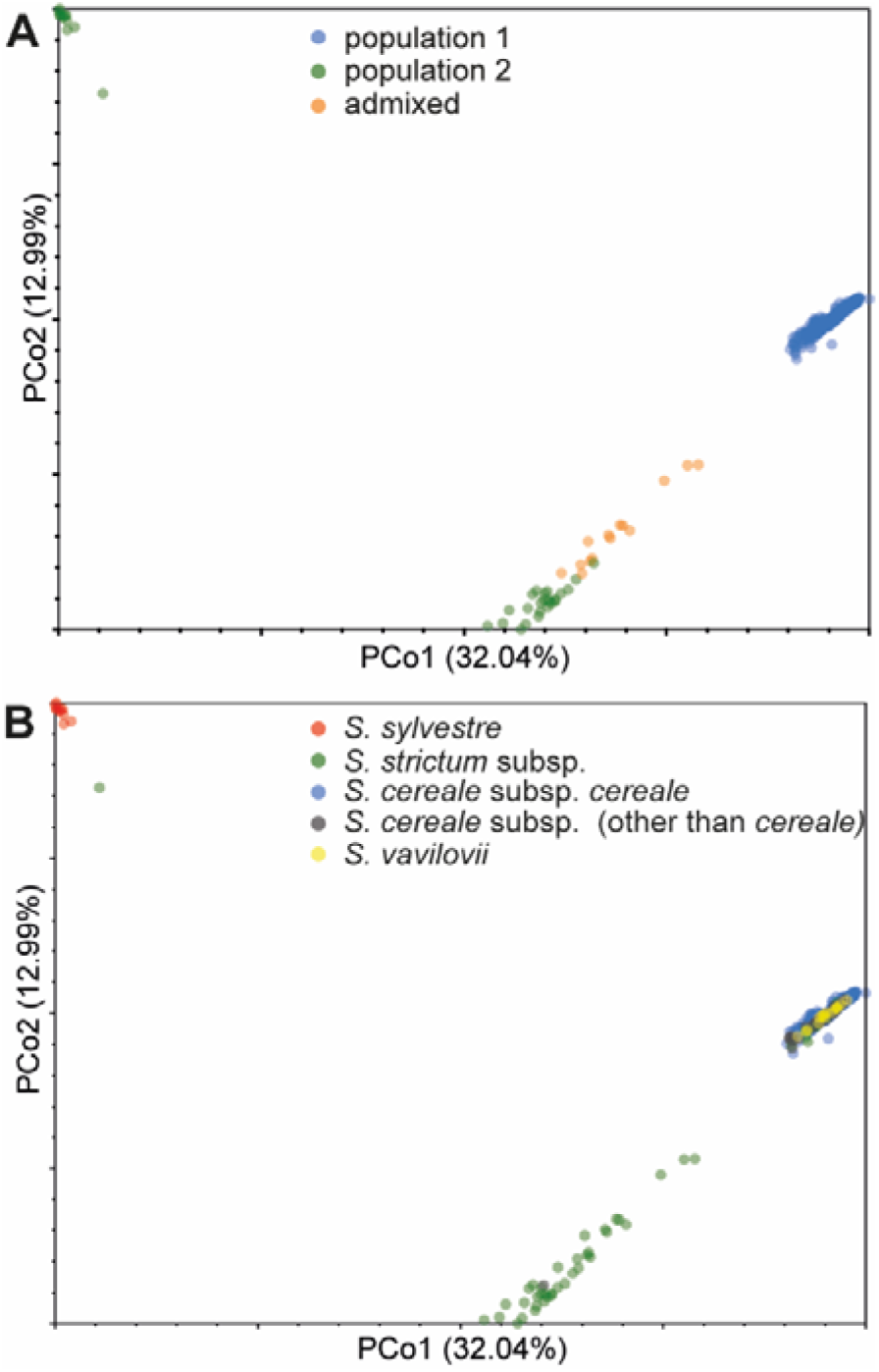
Principal Coordinates Analysis plot showing relationships between 478 rye accessions genotyped with 12 846 SNPs. **A** Accessions labelled according to STRUCTURE-based population assignments. **B** accessions labelled according to taxonomy.

### NJ clustering

Three major clusters could be distinguished in the NJ tree showing phylogenetic relationships between accessions: A1, A2, and A3 (Figure 2, Table S1). Cluster A3 could be further subdivided into four subclusters: A3.1-A3.4. The clustering was in very good agreement with the STRUCTURE and PCoA results. Accessions from population 2 were grouped in clusters A1 and A2 in the NJ tree, corresponding to two smaller groups of accessions visible in the PCoA plot (Figure S4). Accessions assigned to population 1 formed cluster A3. Admixtures were placed in the outer region of cluster A2, adjacent to cluster A3.1 (Figure 2A).

**Figure 2.**
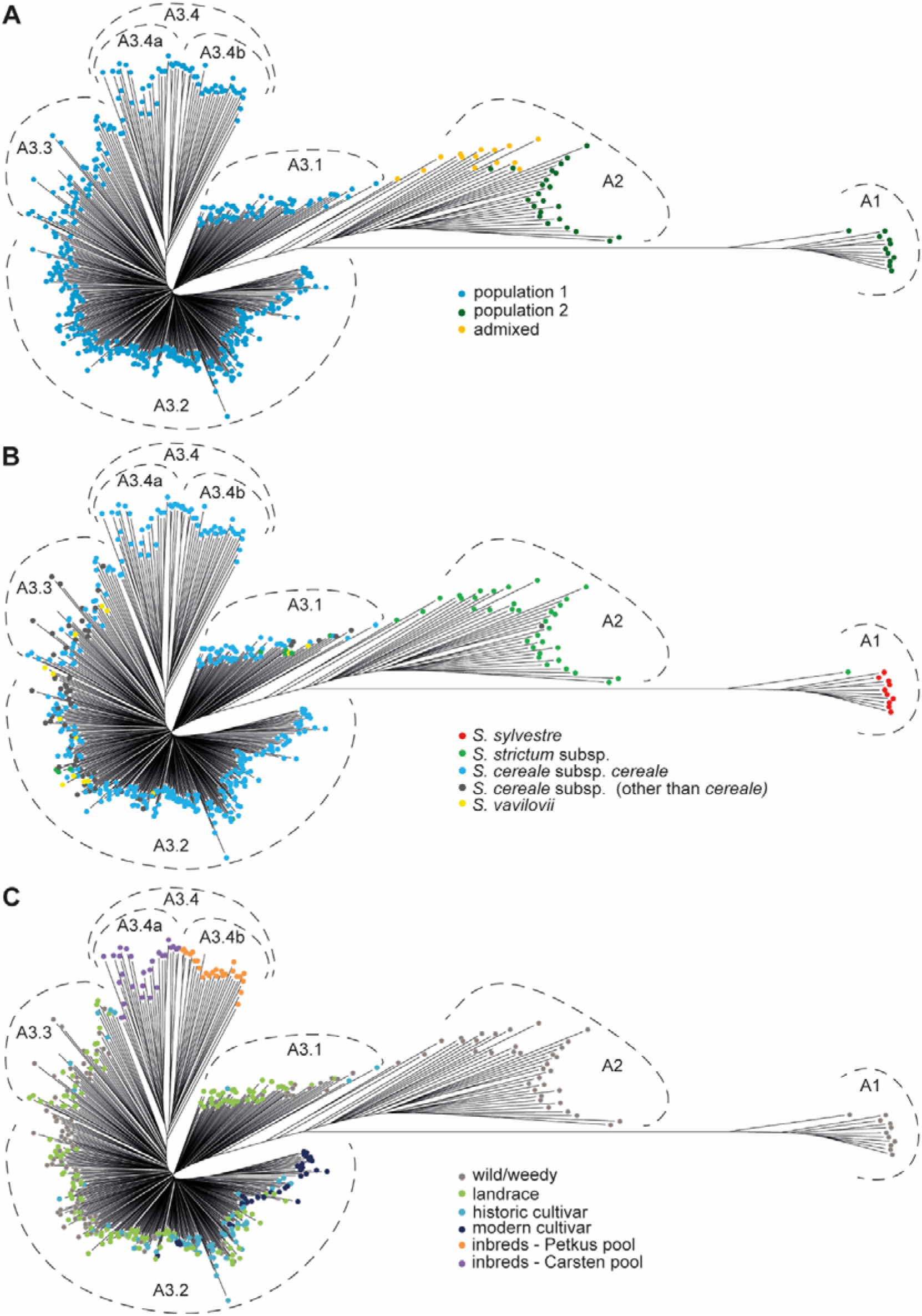
Neighbor-joining tree based on 12 846 SNP markers showing relationships between 478 rye accessions. **A** Accessions labelled according to STRUCTURE-based population assignments NJ trees. **B** accessions labelled according to taxonomy. **C** Accessions labelled according to improvement status.

Clustering corresponded largely with the taxonomy, too (Figure 2B). The most divergent cluster (A1) was composed of *S. sylvestre* accessions, and cluster A2 contained predominantly *S. strictum* accessions, specifically all *S. strictum* subsp. *kuprijanovii* analysed. Cluster A3 contained all cultivated rye *S. cereale* subsp. *cereale* and *S. vavilovii* accessions and almost all wild/weedy *S. cereale* accessions. Wild/weedy S. *cereale* accessions and *S. vavilovii* accessions were dispersed in all A3 subclusters, with exception of subcluster A3.4.

Improvement status of *S. cereale* subsp. *cereale* influenced the clustering (Figure 2C). Inbred lines from a hybrid breeding program formed a separate group (subcluster A3.4), which was divided into two parts corresponding to heterotic pools Carsten and Petkus (A3.4a and A3.4b, respectively). All modern varieties and almost all historical varieties (63 varieties, 87.5 %) were located in cluster A3.2. Historical varieties found in other clusters (A3.1 and A3.3) originated mostly from North America. Apart from varieties, cluster A3.2 contained also 92 landraces (58.2 % of landraces analysed in this study). Landraces occurred also in clusters A3.1 and A.3. (25,3 and 16.5 % of landraces analysed, respectively). The clustering of landraces did not correspond strongly with geographic origin. Cluster A3.2 contained the majority of European (including all landraces from the Balkan region and Southern Europe) and Asian landraces analysed and 31 % of landraces from the Middle East. Landraces from the Middle East were the largest regional germplasm set included in the study (55 accessions from Turkey and Iran, obtained predominantly from the NSGC genebank (36 accessions), but also from PGRC, PAS BG and IPK – 9, 6, and 4 accessions, respectively). A subset of these landraces (ca. 62 % of landraces from the Middle East, mostly Turkish, representing all four genebanks mentioned above) was clearly divergent from the rest and constituted the majority of cluster A3.1. Cluster A3.3 included landraces of various geographic origins: Northern Europe (Finnish and Norwegian), Eastern Europe (Russian), Asia (Afganistan), Western Europe (Germany, Austria and Switzerland) and the Middle East. These landraces originated almost exclusively from IPK and NordGen genebanks.

The highest numbers of accessions for this study were obtained from the following four genebanks: IPK (113), NSGC (97), NordGen (56), and PAS BG (52). Accessions obtained from the IPK genebank covered a broad spectrum of diversity within S. *cereale*/*S. vavilovii* group (Cluster A3) and were dispersed in subclusters A3.1, A3.2 and A3.3, similarly to the accessions from NSGC (Figure S5, Table S1). Accessions from PAS BG were not represented in cluster A3.3 (with exception of a single *S. c*. subsp. *ancestrale* accession), while accessions from NordGen were absent from cluster A3.1. Taken together the accessions obtained for the study provided a good representation of diversity within the *Secale* genus.

### AMOVA

A very high degree of differentiation was found between the two subpopulations indicated by STRUCTURE (F_ST_ = 0.468, 53% percent of total molecular variance attributed to variation within populations, P < 0.001). A very high degree of differentiation was also found for the three accessions groups indicated by PCoA and NJ clustering (corresponding to the A1, A2, and A3 clusters in the NJ tree), with the proportion of molecular variance explained by the differences among populations equal 55% and pairwise population F_ST_ values ranging from 0.428 between accessions groups A2 and A3 to 0.734 between accession groups A1 and A3 (P < 0.001). AMOVA analysis of the six germplasm groups defined based on NJ clustering: A1, A2, A3.1, A3.2, A3.3, and A3.4 attributed 31 % of the total molecular variance to the differences among populations and 69 % to the differences within populations. There was a very high degree of differentiation between groups A1 and A2 and the remaining germplasm groups, with pairwise F_ST_ values ranging from 0.785 to 0.603 and 0.603 to 0.318, respectively. In the remaining germplasm group pairs the degree of differentiation was moderate - pairwise F_ST_ values ranged from 0.054 (between groups A3.1 and A3.2) to 0.205 (between groups A3.2 and A3.4), with the exception of population pair A3.1 and A3.3, where the differentiation was low - pairwise F_ST_=0.035 (P < 0.001, Table S4). There was a moderate degree of differentiation F_ST_=0.092, P < 0.001) between the two groups of lines from hybrid breeding program (A3.4a and A3.4b).

### Diversity indices

Rye accessions were assigned to groups based on improvement status, taxonomy, results of population structure and phylogenetics analyses. Summary of SNP marker numbers, He (expected heterozygosity and Ho (observed heterozygosity) values, and physical map length by germplasm group is given in Table 1, while the information on accessions’ membership in the defined groups is given in Table S1. The numbers of mapped HQ SNPs differentiating the defined groups varied from 10 364 (97.7 % of the HQ, mapped SNPs polymorphic in the whole set) in the *S. strictum* group to only 803 markers (7.6%) in the *S. sylvestre* group. A similar pattern of the chromosomal distribution of SNPs could be observed in most germplasm groups, with the highest proportion of SNPs on chromosomes 2R and 5R and the lowest on chromosomes 1R, 4R and 6R (Figure S6). When accessions were grouped according to taxonomy, a deviation from this pattern was noticeable in the *S. sylvestre* group, where a relatively low proportion of SNPs originated from 5R. In *S. sylvestre* and also in *S. strictum* group a relatively high proportion of SNPs was mapped on 6R. When accession were grouped according to improvement status, a relatively low percentage of SNPs was observed on 5R and relatively high – on 6R in the wild/weedy group.

**Table 1.** Summary of SNP marker numbers, He and Ho values, and physical map length by germplasm group.

| Germplasm group | N | No. of<br>HQ SNPs | No. of<br>mapped<br>HQ SNPs | Proportion<br>of mapped<br>HQ SNPs (%) | mean<br>He | mean<br>Ho | Map<br>length<br>(Mbp) |
| --- | --- | --- | --- | --- | --- | --- | --- |
| All | 478 | 12846 | 10607 | 100 | 0.211 | 0.228 | 6723.25 |
| **Improvement status** |  |  |  |  |  |  |  |
| wild_weedy | 132 | 12837 | 10599 | 99.9 | 0.282 | 0.225 | 6723.25 |
| landrace | 158 | 9181 | 7544 | 71.1 | 0.255 | 0.366 | 6719.24 |
| historic_cultivar | 75 | 7645 | 6194 | 58.4 | 0.279 | 0.466 | 6719.05 |
| modern_cultivar | 36 | 6006 | 4832 | 45.6 | 0.298 | 0.431 | 6713.26 |
| breeding | 53 | 6794 | 5515 | 52.0 | 0.210 | 0.086 | 6716.18 |
| Petkus_pool | 26 | 4280 | 3420 | 32.2 | 0.266 | 0.036 | 6703.50 |
| Carsten_pool | 27 | 4638 | 3698 | 34.9 | 0.269 | 0.127 | 6711.25 |
| **Taxonomy** |  |  |  |  |  |  |  |
| *S. cereale* | 403 | 10450 | 8642 | 81.5 | 0.209 | 0.270 | 6719.24 |
| *S. c. subsp. cereale* | 344 | 9755 | 8031 | 75.7 | 0.220 | 0.287 | 6719.24 |
| *S. c. subsp. "noncereale"* | 59 | 9446 | 7820 | 73.7 | 0.251 | 0.318 | 6719.05 |
| *S. strictum* | 45 | 12562 | 10364 | 97.7 | 0.272 | 0.237 | 6723.25 |
| *S. sylvestre* | 10 | 1045 | 803 | 7.6 | 0.234 | 0.183 | 6622.07 |
| *S. vavilovii* | 14 | 6829 | 5536 | 52.2 | 0.333 | 0.463 | 6713.68 |
| **NJ clusters** |  |  |  |  |  |  |  |
| A1 | 11 | 1602 | 1243 | 11.7 | 0.196 | 0.158 | 6645.31 |
| A2 | 37 | 10181 | 7498 | 70.7 | 0.308 | 0.318 | 6718.51 |
| A3 | 430 | 10482 | 8675 | 81.8 | 0.209 | 0.271 | 6719.70 |
| A3.1 | 59 | 9067 | 7443 | 70.2 | 0.285 | 0.402 | 6719.24 |
| A3.2 | 264 | 9884 | 8160 | 76.9 | 0.223 | 0.328 | 6719.05 |
| A3.3 | 53 | 8088 | 6611 | 62.3 | 0.238 | 0.271 | 6719.51 |
| A3.4 | 54 | 5466 | 4390 | 41.4 | 0.244 | 0.070 | 6715.01 |
| A3.4a | 24 | 4425 | 3537 | 33.3 | 0.272 | 0.145 | 6710.18 |
| A3.4b | 30 | 4603 | 3682 | 34.7 | 0.263 | 0.033 | 6704.52 |
| **Sweep detection sets** |  |  |  |  |  |  |  |
| g1 | 198 | 8952 | 7345 | 69.2 | 0.241 | 0.360 | 6719.05 |
| g2 | 46 | 8594 | 7031 | 66.3 | 0.305 | 0.452 | 6719.24 |
| g3 | 37 | 7640 | 6212 | 58.6 | 0.249 | 0.280 | 6719.51 |
| g4 | 53 | 5447 | 4373 | 41.2 | 0.252 | 0.070 | 6715.01 |
| g4a | 25 | 4524 | 3615 | 34.1 | 0.270 | 0.139 | 6710.18 |
| g4b | 28 | 4511 | 3606 | 34.0 | 0.266 | 0.033 | 6704.52 |

He and Ho values were 0.211 and 0.228 for the whole set. Within established germplasm groups He ranged from 0.196 for the A1 group established based on NJ clustering to 0.333 for the taxonomic group *S. vavilovii* (Table 1). Values of He above 0.3 were also obtained for the group A2 (0.308) and in the sweep detection set g2. The highest Ho values occurred in groups of historic cultivars (0.466) and *S. vavilovii* (0.463). The lowest Ho values, in the range 0.145-0.033, occurred in the germplasm groups containing inbred lines from hybrid breeding program. A low Ho value of 0.183 was also obtained also for the group of *S. sylvestre*, which is a self-pollinating species.

Values of the diversity indices P_S_ (proportion of polymorphic sites), 1 (theta), π (nucleotide diversity), and Tajima D’s values computed for the whole set of 478 accessions and for the established germplasm groups are given in Table S5. When the accession were grouped according to improvement status, the values of diversity indices were the highest in the wild/weedy group, and the lowest in modern cultivars. In taxonomical groups the highest values of diversity indices were observed in *S. strictum* and the lowest in *S. sylvestre*. Average Tajima’s D values were negative in each germplasm group indicating the occurrence of positive selection. The weakest negative Tajima’s D values were recorded in breeding lines. In the cultivated germplasm the strongest negative Tajima’s D value was obtained for modern cultivars, followed by historic cultivars, implying a strong selection in this groups. Among taxonomic groups, the strongest negative Tajima’s D value was noted for *S. vavilovii*.

### Selective sweeps in established groups of cultivated germplasm

Selective sweep detection was performed in groups of cultivated accessions established based on the outcome of NJ clustering (Figure S7, Table S1): g1 (historical and modern cultivars and related landraces from NJ cluster A3.2), g2 (divergent Turkish landraces from cluster A3.1), g3 (diverse landraces from cluster A3.3), and g4 (inbred lines from hybrid breeding program from cluster A3.4), and also, separately g4a (inbred lines from Carsten heterotic pool) and g4b (inbred lines from Petkus heterotic pool). At the adopted settings the algorithms used (SweeD, OmegaPlus, and RAiSD) identified 133 outlier positions in the rye genome in common (Table 2, Table S6). The outliers were located within 13 sweep regions ranging in size from 0.84 Mb to 11.76 Mb (Table S6). The number of sweeps per germplasm group ranged from one (in groups g4a and g4b) to four in group g3. The largest number of outliers (55) was identified in group g4. The putative selective sweeps were dispersed across the rye genome, with the largest number of sweeps (four) detected in chromosome 7R. No sweeps were detected in chromosome 4R (Table 2).

**Table 2.** Chromosomal location of putative sweep regions and numbers of outliers and candidate genes identified in groups of cultivated rye accessions.

| Germplasm group/parameter |  | Chromosome |  |  |  |  |  |  |  | Total |
| --- | --- | --- | --- | --- | --- | --- | --- | --- | --- | --- |
|  |  | 1R | 2R | 3R | 4R | 5R | 6R | 7R | Un |  |
| g1 | no. of sweeps |  |  |  |  | 1 |  | 2 |  | 3 |
|  | no. of outliers |  |  |  |  | 1 |  | 2 (1+1)<sup>a</sup> |  | 3 |
|  | no. of candidate genes |  |  |  |  | 4 |  | 21 (10+11)<sup>a</sup> |  | 25 |
| g2 | no. of sweeps | 1 |  |  |  |  |  | 1 |  | 2 |
|  | no. of outliers | 8 |  |  |  |  |  | 1 |  | 9 |
|  | no. of candidate genes | 6 |  |  |  |  |  | 14 |  | 20 |
| g3 | no. of sweeps |  | 1 | 1 |  | 1 |  |  | 1 | 4 |
|  | no. of outliers |  | 1 | 8 |  | 2 |  |  | 1 | 12 |
|  | no. of candidate genes |  | 9 | 10 |  | 26 |  |  | 0 | 45 |
| g4 | no. of sweeps |  |  |  |  |  | 1 | 1 |  | 2 |
|  | no. of outliers |  |  |  |  |  | 23 | 29 |  | 52 |
|  | no. of candidate genes |  |  |  |  |  | 23 | 28 |  | 51 |
| g4a | no. of sweeps |  |  |  |  | 1 |  |  |  | 1 |
|  | no. of outliers |  |  |  |  | 13 |  |  |  | 13 |
|  | no. of candidate genes |  |  |  |  | 15 |  |  |  | 15 |
| g4b | no. of sweeps | 1 |  |  |  |  |  |  |  | 1 |
|  | no. of outliers | 44 |  |  |  |  |  |  |  | 44 |
|  | no. of candidate genes | 14 |  |  |  |  |  |  |  | 14 |
| Total | no. of sweeps | 2 | 1 | 1 | 0 | 3 | 1 | 4 | 1 | 13 |
|  | no. of outliers | 52 | 1 | 8 | 0 | 16 | 23 | 32 | 1 | 133 |
|  | no. of candidate genes | 20 | 9 | 10 | 0 | 45 | 23 | 63 | 0 | 170 |

<sup>a</sup> numbers in brackets indicate the number of outliers/candidate genes in each of the respective sweeps.

### Candidate genes in the putative selective sweep regions

In total, 170 putative candidate genes were found in the Lo7 genome in the vicinity of the outlier positions within the identified sweeps (Table S7). The number of candidate genes per chromosome ranged from 63 on chromosome 7R to nine on chromosome R2. Within germplasm groups the highest number of candidate genes was identified in g4 (inbred lines from hybrid breeding program) – 51 genes, followed by g3 (divergent Turkish landraces) – 45 genes. Within the candidate genes, we identified the ones located in the vicinity of the outlier/s with the highest values of the statistics computed by the sweep detection algorithms. These candidate genes are listed in Table 3.

**Table 3.**
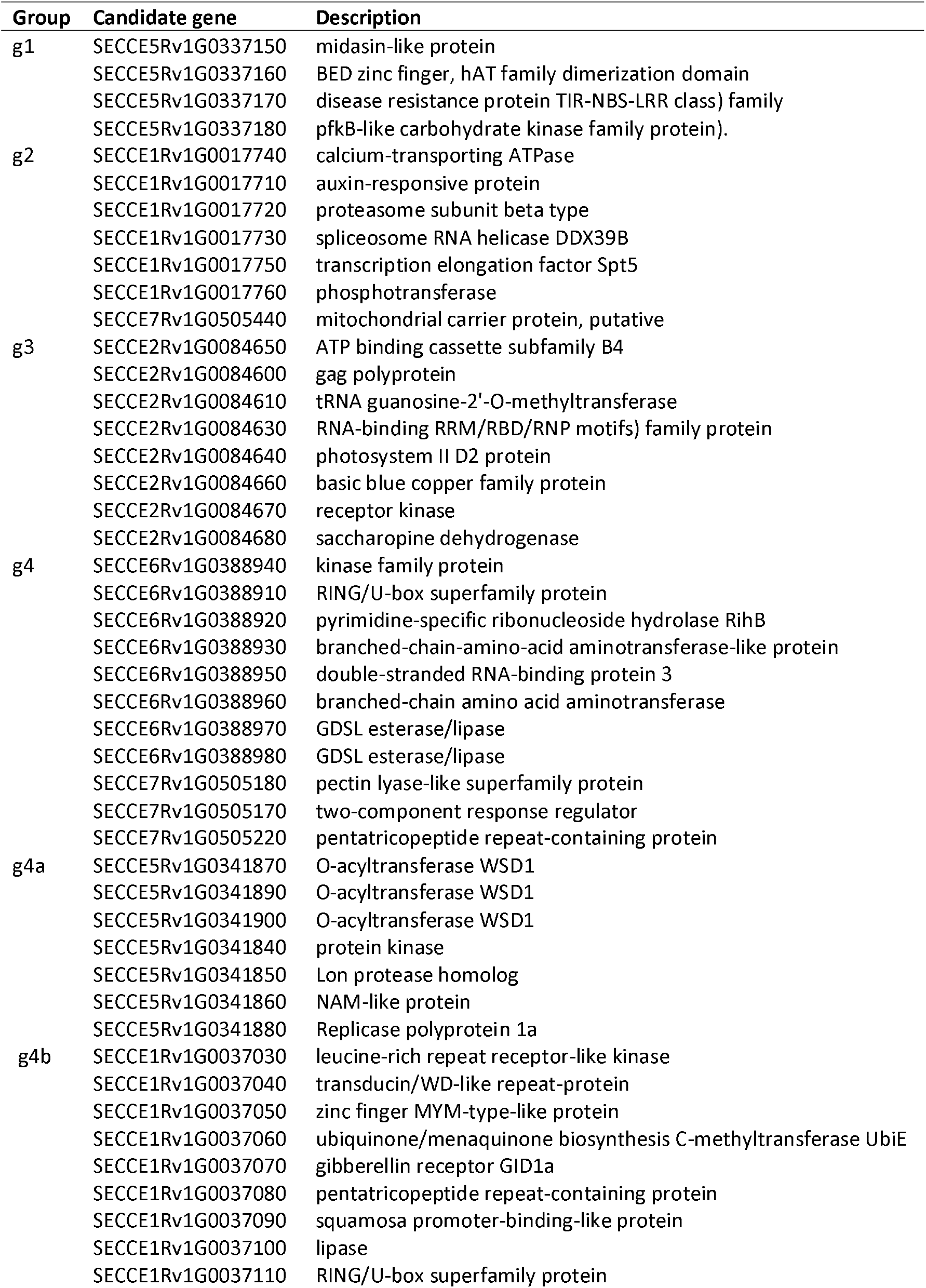
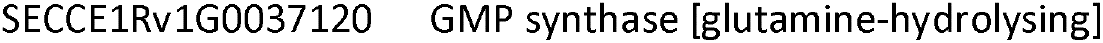
Candidate genes located in the vicinity of the outliers with the highest values of the statistics computed by the sweep detection algorithms.

Gene ontology (GO) enrichment analysis, performed on the identified putative candidate genes, revealed, that 34 GO terms were significantly overrepresented (Table S8), including glycerolipid biosynthetic process, diacylglycerol O-acyltransferase activity, phosphorelay signal transduction system, polygalacturonase activity, embryo sac morphogenesis, and pollen sperm cell differentiation.

### Correspondence to previously identified domestication genes

Based on the literature we compiled a list of several cereal domestication/improvement genes and using BLAST [31] determined the location of their putative homologues in the Lo7 genome (Table S9). We found that those genes were located outside of selective sweeps identified in this study. Previously, selective sweep detection was performed in rye based on the Weining genome sequence by [17]. We located in the genome sequence of Lo7 sequences homologous to putative candidate genes identified by Li et al. and found, that several of them were located in the vicinity (less than 5 Mb) of outliers indicated in this study. This coinciding location was found for the following five candidate genes identified by Li et al.: ScWN1R01G158700_LOC_Os01g08320 (auxin and brassinosteroid hormone responses and plant morphogenesis), ScWN2R01G091200_LOC_Os07g47670 (hypoxia signalling, Pi uptake and accumulation), ScWN2R01G169300_LOC_Os10g25130 (regulation of starch storage in endosperm, internode elongation, domestication traits), ScWN5R01G313900_LOC_Os08g41880 (phosphate deficiency adaptations), ScWN7R01G263700_LOC_Os08g44400 (disease resistance, stress response). On the other hands, while many of the putative candidate genes identified in this study represented the same gene families as the recently described putative candidate genes targeted by selection in rye [32] - for example lipase, gibberellin receptor GID1a, pentatricopeptide repeat-containing protein, leucine-rich repeat protein kinase family protein - their genomic locations did not overlap.

## DISCUSSION

### Genetic diversity within *Secale* genus and within cultivated ryes

The first aim of this study was a detailed analysis of genetic diversity structure in a broad collection of diverse rye germplasm and the identification of germplasm groups suitable for detection of selective sweeps. Several studies on rye genetic diversity were carried out to date, using SSR, array-based (DArT), GBS, and, recently, whole genome resequencing data [18, 19, 21, 28, 32, 33]. In the previous genome-wide studies deploying high-density genotyping to analyse rye genetic diversity [16, 19, 21, 32] up to 143 accessions were used. The present study involved the largest number of rye accessions to date (478) and possibly the most diverse, yet balanced germplasm set, ranging from wild species, random mating populations, and hybrids to inbred lines used in hybrid breeding, with wild accessions, landraces, and cultivars/breeding lines each representing ca. one third of the set. To ensure the best possible representation of genetic diversity the accessions were obtained from multiple genebanks and breeding companies and selected to cover a broad spectrum of geographic origins (Table S1). The accessions derived from genebanks IPK and NSGC turned out to largely overlap with respect to their diversity, while some areas of rye diversity space were not represented by accessions derived from PAS BG and NordGen. However, the number of accessions derived from these genebanks and sampled in this study is too small to justify a suggestion that there are gaps in their rye germplasm collections.

For the detection of SNP variation, the DArTseq genotyping-by-sequencing method was used. This method was previously shown to efficiently target low copy regions of the very large - ca. 8 Gb [16], and highly repetitive (> 90 % [17]) rye genome [34]. Similarly, most of DArTseq were found to align to intragenic regions in wheat [35]. DArTseq genotyping proved to be a suitable tool for high-density genome-wide genetic diversity studies and for the detection of selection signals [35–37].

The HQ SNPs identified in this study for the analysis of genetic diversity and population structure provided good coverage of the rye genome, spanning ca. 99.7% of the reference genome assembly, with similar proportion of markers originating from individual chromosomes (between ca. 11 and 17 %, Table S3). Similarly like in other cereal species, such us barley [38], and wheat [35], a larger proportion of HQ SNP markers segregated in the wild thn in cultivated rye (*S. cereale* subsp. *cereale*) accessions (over 99.9% vs. 75.7%, Table 1), in consistence with the assumption that domestication and improvement resulted in a decrease in diversity, and that crop wild relatives are a treasure trove of untapped and potentially valuable variation for crop improvement [39, 40].

STRUCTURE analysis suggested the presence of two subpopulation in the analysed collection (K=2), dividing the set in to two groups - the first consisting of *S. sylvestre* and *S. strictum* accessions and the second containing *S. vavilovii* and *S. cereale* (both cultivated and weedy) accessions. However, the implementation of Delta K method to the identification of the number cluster explain best the structure within the analysed germplasm set often indicates K=2 as the highest level of hierarchical structure, even if the structure is more complex. Hence, the use of other methods in conjunction with Delta K is recommended [41–43]. Therefore, we further examined relationships between accessions using PCoA and NJ clustering. The results of both analyses were in very good agreement and indicated a more complex structure within *Secale* genus – three main complexes corresponding to the taxonomy: the most divergent *S. sylvestre* complex, the *S. strictum* complex, and the *S. vavilovii*/*S. cereale* complex. This outcome agrees with results of previous studies on rye genetic diversity [18, 19, 21], and is also well supported by the outcome of AMOVA analysis and pairwise F_ST_ values indicating a very high degree of differentiation between these three germplasm groups.

A large number of *S. strictum* subsp. samples included in the study (45 accessions, which originated from different genebanks) allowed us to gather novel information concerning its genetic diversity. We revealed a considerable genetic diversity of this genus, as demonstrated by high values of genetic diversity indices and the results of STRUCTURE analysis, with its representatives present in both populations indicated and also classified as admixtures. NJ clustering indicated differences between *S. strictum* subspecies analysed. *S. stricum* subsp. *kuprijanovii* (14 accessions) turned out to be the most homogenous group, located exclusively in cluster A2, while accessions of *S. strictum* subsp. *anatolicum* and *S. stricum* subsp. *strictum* (eight and 21 accessions, respectively) occurred, besides cluster A2, also in clusters A3.1 and A3.2, intermixed with cultivated and weedy accessions of *Secale cereale*. However, it cannot be excluded, that some of this *S. strictum* samples placed within *S. cereale* complex are only erroneously described as *S. strictum*, since a morphological description was not performed within this study. We find that evaluation of morphological characters would be advisable as a part of future molecular studies on rye taxonomy and phylogeny to exclude possible misclassification of some accessions. On the other hand, the diversity of ten *S. sylvestre* accessions analysed in this turned out to be very small, indicating a need for a follow-up, more detailed examination, to ensure proper safeguarding of the genetic potential of this wild relative of cultivated rye. In consistence with the earlier molecular reports examining taxonomic relationships within the genus *Secale* [18, 21, 32], all the 14 *S. vavilovii* accessions analysed were intermixed with the *S. cereale* accessions in clusters A3.1, A3.2 and A3.3 of the NJ tree. Referring to the explanation of Zohary at al. [22] on the controversies regarding the taxonomic position of *S. vavilovii*, this result would suggest that the samples analysed in this study were “true” *S. vavilovii* forms. Surprisingly, we observed a strong negative Tajima’s D value in this germplasm group which could be a result of an unintentional selection during genebank conservation, for example caused by a low germination rate during regeneration. A coanalysis with *S. vavilovii* samples from a recent collection would be needed to verify this hypothesis. Based on the analysis of our large germplasm set, comprising 58 wild/weedy *Secale cereale* accessions (*S. c*. subsp. *afghanicum, S. c*. subsp. *ancestrale, S. c*. subsp. *dighoricum, S. c*. subsp. *rigidum*, and *S. c*. subsp. *segetale*) we were able to confirm the lack of separation between weedy and cultivated forms of *Secale cereale*, indicated by earlier studies [19, 21, 32] and suggesting a strong gene flow between these two groups.

We detected the presence of genetic clusters within cultivated ryes and the influence of improvement status on the clustering. NJ clustering indicated a genetic distinctiveness of inbreds used in rye hybrid breeding from the remaining cultivated rye accessions, with the two heterotic pools forming separate clusters in the NJ tree. However, the degree of genetic differentiation between heterotic pools, measured by F_ST_ value was moderate. Further, the remaining cultivated accessions formed three major clusters, one containing the majority of cultivars and related landraces, and two additional clusters comprising mostly rye landraces. Thus, the study confirmed the indication from earlier works [18, 27, 28], that a large portion of the genetic diversity of rye landraces is not represented in rye cultivars, especially in the modern ones. Similar patterns of genetic diversity distribution between landraces and cultivars, reflecting strong selection for several key traits, which occurs during breeding/and or initial, historic choices of germplasm for breeding programs, were reported also for other crops [35, 44]. In this study we identified a very distinct group of rye landraces (cluster A3.1) mostly from Turkey. This geographic region is important for rye evolution as the probable area of origin of cultivated rye and the main centre of diversity [22, 25]. Therefore, we postulate that this germplasm group should be of special interest in conservation efforts, and also future allele-mining projects aimed at identification of novel variation for rye improvement [27].

### Detection of genome regions targeted by selection

Detection of selection targets was attempted in rye for the first time by Bauer at al. [45] who used the F_ST_ outlier approach and the *X*^*T*^*X* statistic in pairwise comparisons between three germplasm groups: 38 and 46 inbred lines representing the seed and pollen parent pools used in rye hybrid breeding, respectively, and 46 individuals representing rye genetic resources. The analyses were done based on genotypic data obtained with the use of Rye600k array. In each comparison numerous outlier markers were identified, which clustered in a few distinct genome regions, and in total 27 putative selection targets – rye orthologes of cloned and functionally characterised rice genes – were found in these regions. Functions of these putative orthologues were related, among others, to plant height, grain size and number, pollen germination ability, other plant development and morphology functions, abiotic and biotic stress, and regulation of various physiological processes. Subsequently, Li et al. [17] identified loci potentially involved in the domestication of rye based on GBS data of 101 accessions reported by Schreiber et al. [19]. Specifically, SNPs differentiating 81 cultivated rye and five *S. vavilovii* accessions and three selective sweep detection methods (reduction of diversity (ROD), genome-wide scan of fixation index (F_ST_) and cross-population composite likelihood ratio (CP-CLR)) were used. As a result, 11 selective sweeps with the total of three candidate rye genes, related to brassinosteroid signalling, the transition from vegetative to floral development, and also to tillering and grain yield regulation, were detected in common by the three approaches. The number of candidate genes detected by at least one tool ranged from 10 to 21. Recently, based on resequencing data, Sun et al. [32] performed identification of genes targeted by selection during domestication using F_ST_, XP CLR and ROD approaches on cultivated and weedy ryes from a worldwide set of 116 accessions. As a result selective sweeps with 279 candidate genes were identified by at least two methods, among them, genes related to plant height, disease resistance, tiller number and grain yield, and also genes with shattering-related functions.

In this study, we used a different approach to detect genome regions targeted by positive selection in rye. Each of the defined rye germplasm groups was analysed separately with three algorithms detecting various distinct signatures of selective pressure: SweeD [46], OmegaPlus [47], and RAiSD [48]. Previously, these algorithms were used successfully to identify selective sweeps signals in various plan germplasm sets, such as African rice, maize inbred lines adapted to African highlands, Canadian spring wheat cultivars, and wild strawberries [36, 41, 49, 50]. We performed sweep detection in cultivated accessions from each of the identified clusters within *S. cereale* complex. To reduce the length of the manuscript and to focus on the most probable selection targets, we reported only the sweeps that were identified in common by the three algorithms used and listed the potential candidate genes located in the vicinity of the detected outliers.

Genome-wide studies on the influence of selection on the crop plant genome demonstrated that numerous loci, scattered across the genome, are targeted by selection pressure [8, 12]. In accordance with those findings, we detected a total of 133 outliers that were located in 13 putative sweeps, dispersed in the rye genome. The lengths of selective sweeps regions identified in this study ranged from 0.84 to 11.76 Mb and were comparable to those reported in rye by [17] (2 – 37 Mb) and [32] (0.11-11 Mb).

We were not able to find a clear correspondence between positions of outliers located in this study and the location of known cereal domestication genes orthologs in the rye genome. However, this result is not altogether surprising since the methods used in this study are dedicated to detection of recent and strong positive selection [9]. The overlap between candidate genes for selection reported previously in rye [17, 32] and those identified in this study was very small. Lack of consistency in the results of selective sweep detection studies is frequently encountered in literature [36, 50], and is associated with various factors, including differences in the type of molecular markers used, marker density and genome coverage, size and the diversity of the germplasm set analysed, and the sweep detection method applied [50, 51]. To circumvent some of these limitations and to improve the reliability of sweep detection, it has been suggested to use different sweep detection methods in parallel [52]. We followed this route and used three tools detecting different signatures od selective pressure and reported the overlap of the results from the three tools, providing numerous novel candidate genes targeted by selection in cultivated rye for further studies.

### Potential functions of identified candidate genes

Candidate genes under selection identified so far in crop plants often represent functions related to response to environmental stimuli, such as biotic and abiotic stress resistance, plant architecture, seed size and composition [8, 17, 32, 49, 50]. Similarly, many of the putative candidate genes identified in this study are related to the aforementioned traits.

To prioritise the candidates we used the values of the statistics computed by the algorithms SweeD, OmegaPlus and RAiSD, and we also performed GO term enrichment analysis. One of the significantly enriched GO terms was phosphorelay signal transduction system, which is involved in turning on and off cellular responses to environmental stimuli [53, 54], such as light, cold, or drought. GO enrichment analysis also indicated the GO terms glycerolipid biosynthetic process and diacylglycerol O-acyltransferase activity. Among candidate genes, these two GO terms were represented by several O-acyltransferases WSD1. Three of them were located in the vicinity of an outlier with the highest statistic score, which was identified in group g4a (Carsten heterotic pool). O-acyltransferases WSD1 are involved in cuticular wax biosynthesis, which acts in plants as a protective barrier against biotic and abiotic stresses, including drought [55–57]. Genes representing GO term polygalacturonase activity, were also found to be significantly enriched among putative candidates for selection. This group consisted of several pectin lyase-like genes. Multiple biological functions are being attributed to pectin lyases, such as roles as extracellular virulence agents and roles in plant growth and development, including pollen maturation and pollen tube growth [58]. We also observed significant enrichment of GO terms related to plant fertility and reproduction: embryo sac morphogenesis, and pollen sperm cell differentiation. The candidate genes located in the vicinity of most statistically significant outliers included further genes with obvious connections to traits relevant for plant breeding, such as, among others, genes encoding gibberellin-receptor GID1a, GDSL esterases/lipases, pentatricopeptide repeat-containing proteins, TIR-NBS-LRR class disease resistance protein. Gibberellin-receptors GID1 are key elements in gibberellin signal transduction in plants, and therefore play a role in the control of various aspects of plant growth and fertility, including seed germination and biomass production [59, 60]. GDSL esterases/lipases are active during seed germination and play important roles in plant metabolism, growth and development, including seed development. Pentatricopeptide repeat containing proteins are a large family of proteins regulating gene expression at the RNA level [61]. Most of the known *Restorer of Fertility* (*Rf*) genes are members of this family [62]. TIR-NBS-LRR genes constitute one of two groups of the nucleotide-binding leucine-rich repeat (NB-LRR) family, comprising most of the plant pathogen resistance genes [63].

### Conclusions

Based on a genome-wide detailed analysis of genetic diversity structure in a broad collection of diverse rye germplasm, we identified three complexes within the *Secale* genus: *S. sylvestre, S. strictum* and *S. cereale*/*vavilovii*. We revealed a relatively narrow diversity of *S. sylvestre*, very high diversity of *S. strictum*, and signatures of strong positive selection in *S. vavilovii*. Within cultivated ryes we detected the presence of genetic clusters and the influence of improvement status on the clustering. Rye landraces represent a reservoir of variation for breeding, since a large portion of their diversity is not represented in rye cultivars, and especially a distinct group of landraces from Turkey should be of special interest as a source of untapped variation. Using three sweep detection algorithms we found that signatures of selection are dispersed in the rye genome and identified 170 putative candidate genes targeted by selection in cultivated rye related, among others, to response to various environmental stimuli (such as pathogens, drought, cold), plant fertility and reproduction (pollen sperm cell differentiation, pollen maturation, pollen tube growth), plant growth and biomass production. Our study provides useful information for efficient management of rye germplasm collections, that can help to ensure proper safeguarding of their genetic potential and provides numerous novel candidate genes targeted by selection in cultivated rye for further functional characterisation and allelic diversity studies.

## METHODS

### Plant material

In total of 478 rye (*Secale* sp.) accessions representing different geographic origins and improvement status belonging to 17 taxonomic units were analysed: 134 wild or weedy accessions, 161 landraces, 75 historic cultivars, 36 modern cultivars, 58 breeding lines and 14 accessions of an unknown improvement status. The germplasm collection used was composed of three sets: (i) germplasm set 1 consisted of 340 accessions obtained as seed samples from several genebanks and breeding companies, (ii) germplasm set 2 consisted of 54 rye inbred lines representing heterotic pools used in rye hybrid breeding at HYBRO Saatzucht [64], (iii) germplasm set 3 consisted of 84 accessions described in [21]. Information on accessions used in the study is listed in Table S1.

### DNA isolation and genotyping

For accessions from germplasm set 1 tissue was collected and DNA isolated as described in [27]. Information on DNA isolation for samples from germplasm sets 2 and 3 can be found, respectively, in [64] and [21]. GBS genotyping (DArTseq) was performed at Diversity Array Technology Pty Ltd., Bruce ACT, Australia (http://diversityarrays.com) as described in [65]. For selection of high quality (HQ) SNPs for population structure and genetic diversity analyses the following thresholds were adopted: reproducibility >95%, minor allele frequency (MAF) >0,01), and maximum missing data <10%. SNP filtering was done with R package dartR [66]. PIC values were calculated using the formula by [67].

### Population structure and genetic diversity analyses

The number of clusters (K) capturing the major structure in data was identified with STRUCTURE 2.3.4 software [68] using the admixture model and correlated allele frequencies and the following settings: length of the burn-in period 100,000, number of MCMC replications after burn-in: 10,000. For each number of K tested, ranging from 1 to 14, five independent iterations were performed. The Evanno method [43] implemented in the Structure Harvester [69] was used to identify the number of subpopulations (K) explaining best the population structure in the set. Computation of distance matrices was conducted using R packages adegenet [70, 71], stats and dartR [66]. MEGA11 software [72] was used to construct a Neighbor Joining dendrogram. Principal Coordinate Analysis was done using NTSYSpc 2.2 [73]. AMOVA was performed with GenAlEx 6.503 [74, 75]. He and Ho values were calculated for defined germplasm groups using dartR. A variant call format (vcf) file with markers, which aligned to the reference genome and fulfilled the above specified quality criteria, and the corresponding marker scores was converted using TASSEL v5.2.86 [76] to PHILIP interleaved format and used as input for MEGA11 [72] to calculate the number of segregating sites (S), the proportion of polymorphic sites (Ps), Theta (θ), nucleotide diversity (π), and Tajima’s D [77] for each defined germplasm group.

### Selective sweep detection and GO term enrichment analysis

Three algorithms: i) SweeD [46], ii) OmegaPlus [47], and iii) RAiSD [48] were used to detect selective sweeps in defined groups of cultivated rye germplasm. SweeD is based on the Site Frequency Spectrum (SFS), since an increase of high- and low-frequency derived variants is expected in the proximity of a beneficial mutation. OmegaPlus algorithm relies on Linkage Disequilibrium (LS) patterns, where it is assumed that the LD levels remain high at each side of the beneficial mutation, and drop dramatically for loci across the beneficial mutation. RAiSD detects selective sweeps using multiple signatures of selection: changes in the amount of genetic diversity, SFS, and LD, while relying on SNP vectors. The analyses were run as described earlier [36, 49, 78] on the vcf input file of Dataset-1. Only physically mapped SNPs located outside centromere regions were included in the analyses. The top 2% of the highest scores detected by all three algorithms were declared putative selective sweeps. Two or more adjacent, partially overlapping sweep regions were assumed to be one selective sweep. The identified outlier positions within each putative sweep (plus/ minus 500 kb) were used to search for candidate genes in the reference genome of the rye inbred line Lo7 and the accompanying annotation file [16]. To determine if any functional classes were over-represented among the candidate genes GO enrichment analysis was performed using FUNC-E h(ttps://github.com/SystemsGenetics/FUNC-E) with the *P*-value criterion of <0.01.

## Additional files

### Additional file 1

**Figure S1**. Summary of information on 12 846 HQ SNPs polymorphic in 478 diverse rye accessions. **A**. Histogram of polymorphic information content (PIC) values. **B**. Histogram of minor allele frequency (MAF) values.

**Figure S2**. Plot of Delta K values for number of assumed subpopulations (K) ranging from 2 to 14.

**Figure S3**. Population structure of 478 rye accessions at K=2 based on 12 846 SNPs. Each accession is represented by a vertical stripe partitioned into coloured segments with lengths representing the membership fractions in the inferred clusters. The order of accessions in the plot is the same as in the Table S1.

**Figure S4**. Principal Coordinates Analysis plot showing relationships between 478 rye accessions genotyped with 12846 SNPs with accessions labelled according to the outcome of NJ clustering.

**Figure S5**. Neighbor-joining tree based on 12 846 SNP markers showing relationships between 478 rye accessions with accession labelled according to source.

**Figure S6**. Chromosomal distribution of polymorphic SNPs by germplasm group. **A**. Accessions grouped according to improvement status. **B**. Accessions grouped according to taxonomy. **C**. Accessions grouped according to the outcome of NJ clustering. **D**. Accessions grouped according to their membership in sweep detection sets.

**Figure S7**. Neighbor-joining tree based on 12 846 SNP markers showing relationships between 478 rye accessions with branch colour indicating cultivated rye accessions belonging to the respective sweep detection set.

### Additional file 2

**Table S1**. Information on rye accessions used in the study, including genebank accession number, name, source, taxon, improvement status, country of origin, geographic region, group memberships based on STRUCTURE analysis and NJ clustering, and sweep detection set membership.

**Table S2**. List of 12486 HQ DArTseq markers used in this study, including their sequences and position in the Lo7 reference genome.

**Table S3**. Chromosomal distribution of SNPs by germplasm group.

**Table S4**. Pairwise population F_ST_ values for accessions groups based on NJ clustering.

**Table S5**. Values of genetic diversity indices for the collection of 478 rye accessions and each established germplasm group.

**Table S6** Information on common outliers and sweeps detected by all three methods (SweeD, OmegaPlus, and RAiSD) in groups of cultivated rye accessions.

**Table S7**. List of putative candidate genes targeted by selection in cultivated rye

**Table S8**. Enriched GO terms for putative candidate genes from the selective sweep regions.

**Table S9**. Literature based list of known cereal domestication/improvement genes and locations of their putative homologues in the Lo7 genome.

## DECLARATIONS

### Ethics approval and consent to participate

Not applicable

### Consent for publication

Not applicable

### Availability of data and materials

Data generated or analysed during this study are included in this published article and its supplementary information files or are available from the corresponding author on reasonable request.

### Authors’ contributions

AH, EB, KT, HBB performed the experiments. AH, EB, NA, LB, PG, HBB analysed the data. NA, LB, BH, DS, RD, MS contributed data/analysis tools. HBB conceived, designed and supervised the experiments, wrote the manuscript. AH, NA, BH, DS, RD, MS, HBB reviewed and edited the manuscript. All authors read and approved the final manuscript.

## References

1. Doebley JF, Gaut BS, Smith BD. The Molecular Genetics of Crop Domestication. Cell. 2006;127:1309–21.

2. Abbo S, Pinhasi R, Gopher A, Saranga Y, Ofner I, Peleg Z. Plant domestication versus crop evolution1: a conceptual framework for cereals and grain legumes. Trends Plant Sci. 2014;19:351–60.

3. Lin Z, Li X, Shannon LM, Yeh CT, Wang ML, Bai G, et al. Parallel domestication of the Shattering1 genes in cereals. Nat Genet. 2012;44:720–4.

4. Pourkheirandish M, Hensel G, Kilian B, Senthil N, Chen G, Sameri M, et al. Evolution of the grain dispersal system in barley. Cell. 2015;162:527–39.

5. Palaisa K, Morgante M, Tingey S, Rafalski A. Long-range patterns of diversity and linkage disequilibrium surrounding the maize Y1 gene are indicative of an asymmetric selective sweep. Proc Natl Acad Sci U S A. 2004;101:9885–90.

6. Yano M, Katayose Y, Ashikari M, Yamanouchi U, Monna L, Fuse T, et al. Hd1, a major photoperiod sensitivity quantitative trait locus in rice, is closely related to the Arabidopsis flowering time gene CONSTANS. Plant Cell. 2000;12:2473–83.

7. Pearce S, Saville R, Vaughan SP, Chandler PM, Wilhelm EP, Sparks CA, et al. Molecular characterization of Rht-1 dwarfing genes in hexaploid wheat. Plant Physiol. 2011;157:1820–31.

8. Maccaferri M, Harris NS, Twardziok SO, Pasam RK, Gundlach H, Spannagl M, et al. Durum wheat genome highlights past domestication signatures and future improvement targets. Nat Genet. 2019;51:885–95.

9. Pavlidis P, Alachiotis N. A survey of methods and tools to detect recent and strong positive selection. J Biol Res. 2017;:1–17.

10. De Mita S, Thuillet AC, Gay L, Ahmadi N, Manel S, Ronfort J, et al. Detecting selection along environmental gradients: Analysis of eight methods and their effectiveness for outbreeding and selfing populations. Mol Ecol. 2013;22:1383–99.

11. Ayalew H, Sorrells ME, Carver BF, Baenziger PS, Ma XF. Selection signatures across seven decades of hard winter wheat breeding in the Great Plains of the United States. Plant Genome. 2020;13:1–10.

12. Stetter MG, Gates DJ, Mei W, Ross-Ibarra J. How to make a domesticate. Current Biology. 2017;27:R896–900.

13. Purugganan MD, Fuller DQ. The nature of selection during plant domestication. Nature. 2009;457 February:843–8.

14. Rakoczy-Trojanowska M, Bolibok-Brągoszewska H, Myśków B, Dzięgielewska M, Stojałowski S, Grądzielewska A, et al. Genetics and genomics of stress tolerance. In: Rabanus-Wallace Mt, Stein N, editors. The Rye Genome, Compendium of Plant Genomes. Springer, Cham; 2021. p. 213–36.

15. Crespo-Herrera LA, Garkava-Gustavsson L, Åhman I. A systematic review of rye (Secale cereale L.) as a source of resistance to pathogens and pests in wheat (Triticum aestivum L.). Hereditas. 2017;154:14.

16. Rabanus-Wallace TM, Hackauf B, Mascher M, Lux T, Wicker T, Gundlach H, et al. Chromosome-scale genome assembly provides insights into rye biology, evolution and agronomic potential. Nat Genet. 2021;53:564–73.

17. Li G, Wang L, Yang J, He H, Jin H, Li X, et al. A high-quality genome assembly highlights rye genomic characteristics and agronomically important genes. Nat Genet. 2021;53:574–84.

18. Bolibok-Bragoszewska H, Targonska M, Bolibok L, Kilian A, Rakoczy-Trojanowska M. Genome-wide characterization of genetic diversity and population structure in Secale. BMC Plant Biol. 2014;14:184.

19. Schreiber M, Himmelbach A, Börner A, Mascher M. Genetic diversity and relationship between domesticated rye and its wild relatives as revealed through genotyping-by-sequencing. Evol Appl. 2018; February:1–12.

20. Targońska-Karasek M, Bolibok-Brągoszewska H, Oleniecki T, Sharifova S, Kopania M, Rakoczy-Trojanowska M. Verification of taxonomic relationships within the genus Secale (Poaceae: Pooideae: Triticeae) based on multiple molecular methods. Phytotaxa. 2018;383:128.

21. Al-Beyroutiova M, Sabo M, Sleziak P, Dusinsky R, Bircak E, Hauptvogel P, et al. Evolutionary relationships in the genus Secale revealed by DArTseq DNA polymorphism. Plant Syst Evol. 2016;302:1083–91.

22. Zohary D, Hopf M, Weiss E. Domestication of Plants in the Old World: The origin and spread of domesticated plants in Southwest Asia, Europe, and the Mediterranean Basin. Oxford University Press; 2012.

23. Preece C, Livarda A, Christin PA, Wallace M, Martin G, Charles M, et al. How did the domestication of Fertile Crescent grain crops increase their yields? Funct Ecol. 2017;31:387–97.

24. Fuller DQ. Contrasting patterns in crop domestication and domestication rates: Recent archaeobotanical insights from the old world. Ann Bot. 2007;100:903–24.

25. Behre KE. The history of rye cultivation in Europe. Veg Hist Archaeobot. 1992;1:141–56.

26. Tang ZX, Ross K, Ren ZL, Yang ZJ, Zhang HY T C, et al. Secale. In: C K, editor. Wild Crop Relatives: Genomic and Breeding Resources, Cereals. Berlin Heidelberg: Springer; 2011. p. 367–96.

27. Hawliczek A, Bolibok L, Tofil K, Borzęcka E, Jankowicz-Cieślak J, Gawroński P, et al. Deep sampling and pooled amplicon sequencing reveals hidden genic variation in heterogeneous rye accessions. BMC Genomics. 2020;21:845.

28. Targońska M, Bolibok-Brągoszewska H, Rakoczy-Trojanowska M. Assessment of genetic diversity in Secale cereale based on SSR markers. Plant Mol Biol Report. 2016;34:37–51.

29. Monteiro F, Vidigal P, Barros AB, Monteiro A, Oliveira HR, Viegas W. Genetic Distinctiveness of Rye In situ Accessions from Portugal Unveils a New Hotspot of Unexplored Genetic Resources. Front Plant Sci. 2016;7.

30. Sidhu JS, Ramakrishnan SM, Ali S, Bernardo A, Bai G, Abdullah S, et al. Assessing the genetic diversity and characterizing genomic regions conferring Tan Spot resistance in cultivated rye. PLoS One. 2019;14:1–22.

31. Altschul SF, Gish W, Miller W, Myers EW, Lipman DJ. Basic local alignment search tool. J Mol Biol. 1990;215:403–10.

32. Sun Y, Shen E, Hu Y, Wu D, Feng Y, Lao S, et al. Population genomic analysis reveals domestication of cultivated rye from weedy rye. Mol Plant. 2022;15:552–61.

33. Hagenblad J, Oliveira HR, Forsberg NEG, Leino MW. Geographical distribution of genetic diversity in Secale landrace and wild accessions. BMC Plant Biol. 2016;16:1–20.

34. Borzęcka E, Hawliczek-Strulak A, Bolibok L, Gawroński P, Tofil K, Milczarski P, et al. Effective BAC clone anchoring with genotyping-by-sequencing and Diversity Arrays Technology in a large genome cereal rye. Sci Rep. 2018;8:8428.

35. Sansaloni C, Franco J, Santos B, Percival-Alwyn L, Singh S, Petroli C, et al. Diversity analysis of 80,000 wheat accessions reveals consequences and opportunities of selection footprints. Nat Commun. 2020;11:1–12.

36. Ndjiondjop MN, Alachiotis N, Pavlidis P, Goungoulou A, Kpeki SB, Zhao D, et al. Comparisons of molecular diversity indices, selective sweeps and population structure of African rice with its wild progenitor and Asian rice. Theor Appl Genet. 2019;132:1145–58.

37. Allan V, Vetriventhan M, Senthil R, Geetha S, Deshpande S, Rathore A, et al. Genome-Wide DArTSeq Genotyping and Phenotypic Based Assessment of Within and Among Accessions Diversity and Effective Sample Size in the Diverse Sorghum, Pearl Millet, and Pigeonpea Landraces. Front Plant Sci. 2020;11 December:1–20.

38. Milner SG, Jost M, Taketa S, Mazón ER, Himmelbach A, Oppermann M, et al. Genebank genomics highlights the diversity of a global barley collection. Nat Genet. 2019;51:319–26.

39. Zhang H, Mittal N, Leamy LJ, Barazani O, Song BH. Back into the wild—Apply untapped genetic diversity of wild relatives for crop improvement. Evol Appl. 2017;10:5–24.

40. Brozynska M, Furtado A, Henry RJ. Genomics of crop wild relatives: Expanding the gene pool for crop improvement. Plant Biotechnol J. 2016;14:1070–85.

41. Hu Y, Feng C, Yang L, Edger PP, Kang M. Genomic population structure and local adaptation of the wild strawberry Fragaria nilgerrensis. Hortic Res. 2022;9 October 2021.

42. Janes JK, Miller JM, Dupuis JR, Malenfant RM, Gorrell JC, Cullingham CI, et al. The K = 2 conundrum. Mol Ecol. 2017;26:3594–602.

43. Evanno G, Regnaut S, Goudet J. Detecting the number of clusters of individuals using the software STRUCTURE: a simulation study. Mol Ecol. 2005;14:2611–20.

44. Müller T, Schierscher-Viret B, Fossati D, Brabant C, Schori A, Keller B, et al. Unlocking the diversity of genebanks: whole-genome marker analysis of Swiss bread wheat and spelt. Theor Appl Genet. 2018;131:407–16.

45. Bauer E, Schmutzer T, Barilar I, Mascher M, Gundlach H, Martis MM, et al. Towards a whole-genome sequence for rye (Secale cereale L.). Plant J. 2017;89:853–69.

46. Pavlidis P, Živković D, Stamatakis A, Alachiotis N. SweeD: Likelihood-based detection of selective sweeps in thousands of genomes. Mol Biol Evol. 2013;30:2224–34.

47. Alachiotis N, Stamatakis A, Pavlidis P. OmegaPlus: a scalable tool for rapid detection of selective sweeps in whole-genome datasets. Bioinformatics. 2012;28:2274–5.

48. Alachiotis N, Pavlidis P. RAiSD detects positive selection based on multiple signatures of a selective sweep and SNP vectors. Commun Biol. 2018;1.

49. Wegary D, Teklewold A, Prasanna BM, Ertiro BT, Alachiotis N, Negera D, et al. Molecular diversity and selective sweeps in maize inbred lines adapted to African highlands. Sci Rep. 2019; April:1–15.

50. Semagn K, Iqbal M, Alachiotis N, N’Diaye A, Pozniak C, Spaner D. Genetic diversity and selective sweeps in historical and modern Canadian spring wheat cultivars using the 90K SNP array. Sci Rep. 2021;11:1–16.

51. Narum SR, Hess JE. Comparison of FST outlier tests for SNP loci under selection. Mol Ecol Resour. 2011;11 SUPPL. 1:184–94.

52. Vasemägi A, Nilsson J, Primmer CR. Expressed sequence tag-linked microsatellites as a source of gene-associated polymorphisms for detecting signatures of divergent selection in Atlantic salmon (Salmo salar L.). Mol Biol Evol. 2005;22:1067–76.

53. Mira-Rodado V. New insights into multistep-phosphorelay (Msp)/ two-component system (tcs) regulation: Are plants and bacteria that different? Plants. 2019;8:1–21.

54. Oka A, Sakai H, Iwakoshi S. His-Asp phosphorelay signal transduction in higher plants: Receptors and response regulators for cytokinin signaling in Arabidopsis thaliana. Genes Genet Syst. 2002;77:383–91.

55. Li F, Wu X, Lam P, Bird D, Zheng H, Samuels L, et al. Identification of the wax ester synthase/acyl-coenzyme a:diacylglycerol acyltransferase WSD1 required for stem wax ester biosynthesis in Arabidopsis. Plant Physiol. 2008;148:97–107.

56. Xia K, Ou X, Gao C, Tang H, Jia Y, Deng R, et al. OsWS1 involved in cuticular wax biosynthesis is regulated by osa-miR1848. Plant Cell Environ. 2015;38:2662–73.

57. Laskoś K, Myśków B, Dziurka M, Warchoł M, Dziurka K, Juzoń K, et al. Variation between glaucous and non ⍰glaucous near ⍰isogenic lines of rye (Secale cereale L.) under drought stress. Sci Rep. 2022;:1–17.

58. Cao J. The Pectin Lyases in Arabidopsis thaliana: Evolution, Selection and Expression Profiles. PLoS One. 2012;7.

59. Ge W, Steber CM. Positive and negative regulation of seed germination by the Arabidopsis GA hormone receptors, GID1a, b, and c. Plant Direct. 2018;2.

60. Wang X, Li J, Ban L, Wu Y, Wu X, Wang Y, et al. Functional characterization of a gibberellin receptor and its application in alfalfa biomass improvement. Sci Rep. 2017;7 August 2016:1–12.

61. Manna S. An overview of pentatricopeptide repeat proteins and their applications. Biochimie. 2015;113:93–9.

62. Melonek J, Stone JD, Small I. Evolutionary plasticity of restorer-of-fertility-like proteins in rice. Sci Rep. 2016;6 October:1–12.

63. Lee HA, Yeom SI. Plant NB-LRR proteins: Tightly regulated sensors in a complex manner. Brief Funct Genomics. 2015;14:233–42.

64. Siekmann D, Jansen G, Zaar A, Kilian A, Fromme FJ, Hackauf B. A Genome-Wide Association Study Pinpoints Quantitative Trait Genes for Plant Height, Heading Date, Grain Quality, and Yield in Rye (Secale cereale L.). Front Plant Sci. 2021;12 October:1–23.

65. Targońska-Karasek M, Bolibok-Brągoszewska H, Rakoczy-Trojanowska M. DArTseq genotyping reveals high genetic diversity of polish rye inbred lines. Crop Sci. 2017;57:1906–15.

66. Gruber B, Unmack PJ, Berry OF, Georges A. dartR: An R package to facilitate analysis of SNP data generated from reduced representation genome sequencing. Mol Ecol Resour. 2018;18:691–9.

67. Botstein D, White RL, Skolnick M, Davis RW. Construction of a Genetic Linkage Map in Man Using Restriction Fragment Length Polymorphisms. Am J Hum Gen. 1980;32:314–31.

68. Pritchard JK, Stephens M, Donnelly P. Inference of population structure using multilocus genotype data. Genetics. 2000;155:945–59.

69. Earl D a., vonHoldt BM. STRUCTURE HARVESTER: a website and program for visualizing STRUCTURE output and implementing the Evanno method. Conserv Genet Resour. 2011;4:359–61.

70. Jombart T. Adegenet: A R package for the multivariate analysis of genetic markers. Bioinformatics. 2008;24:1403–5.

71. Jombart T, Ahmed I. adegenet 1.3-1: New tools for the analysis of genome-wide SNP data. Bioinformatics. 2011;27:3070–1.

72. Tamura K, Stecher G, Kumar S. MEGA11: Molecular Evolutionary Genetics Analysis Version 11. Mol Biol Evol. 2021;38:3022–7.

73. Rohlf FJ. NTSYSpc: Numerical Taxonomy System, ver. 2.2. 2008.

74. Peakall R, Smouse PE. Genalex 6: genetic analysis in Excel. Population genetic software for teaching and research. Mol Ecol Notes. 2006;6:288–95.

75. Peakall R, Smouse PE. GenAlEx 6.5: genetic analysis in Excel. Population genetic software for teaching and research--an update. Bioinformatics. 2012;28:2537–9.

76. Bradbury PJ, Zhang Z, Kroon DE, Casstevens TM, Ramdoss Y, Buckler ES. TASSEL: Software for association mapping of complex traits in diverse samples. Bioinformatics. 2007;23:2633–5.

77. Tajima F. Statistical method for testing the neutral mutation hypothesis by DNA polymorphism. Genetics. 1989;:585–95.

78. Ji F, Ma Q, Zhang W, Liu J, Feng Y, Zhao P, et al. A genome variation map provides insights into the genetics of walnut adaptation and agronomic traits. Genome Biol. 2021;22:1–22.

