## Additional File 1 for "Selective sweeps identification in distinct groups of cultivated rye (*Secale cereale* L.) germplasm provides potential candidates for crop improvement"

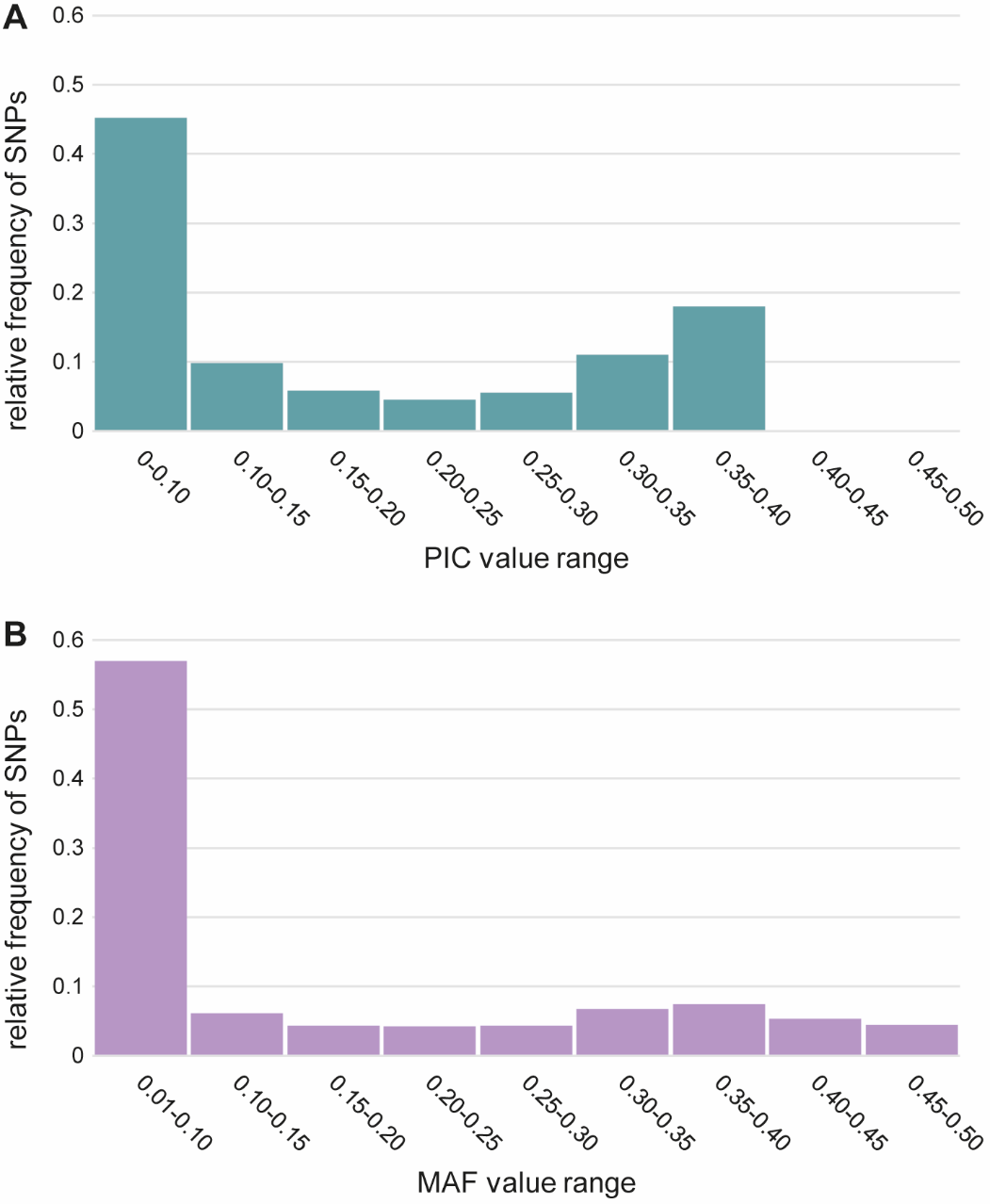

**Figure S1.** Summary of information on 12 846 HQ SNPs polymorphic in 478 diverse rye accessions. **A.** Histogram of polymorphic information content (PIC) values. **B.** Histogram of minor allele frequency (MAF) values.

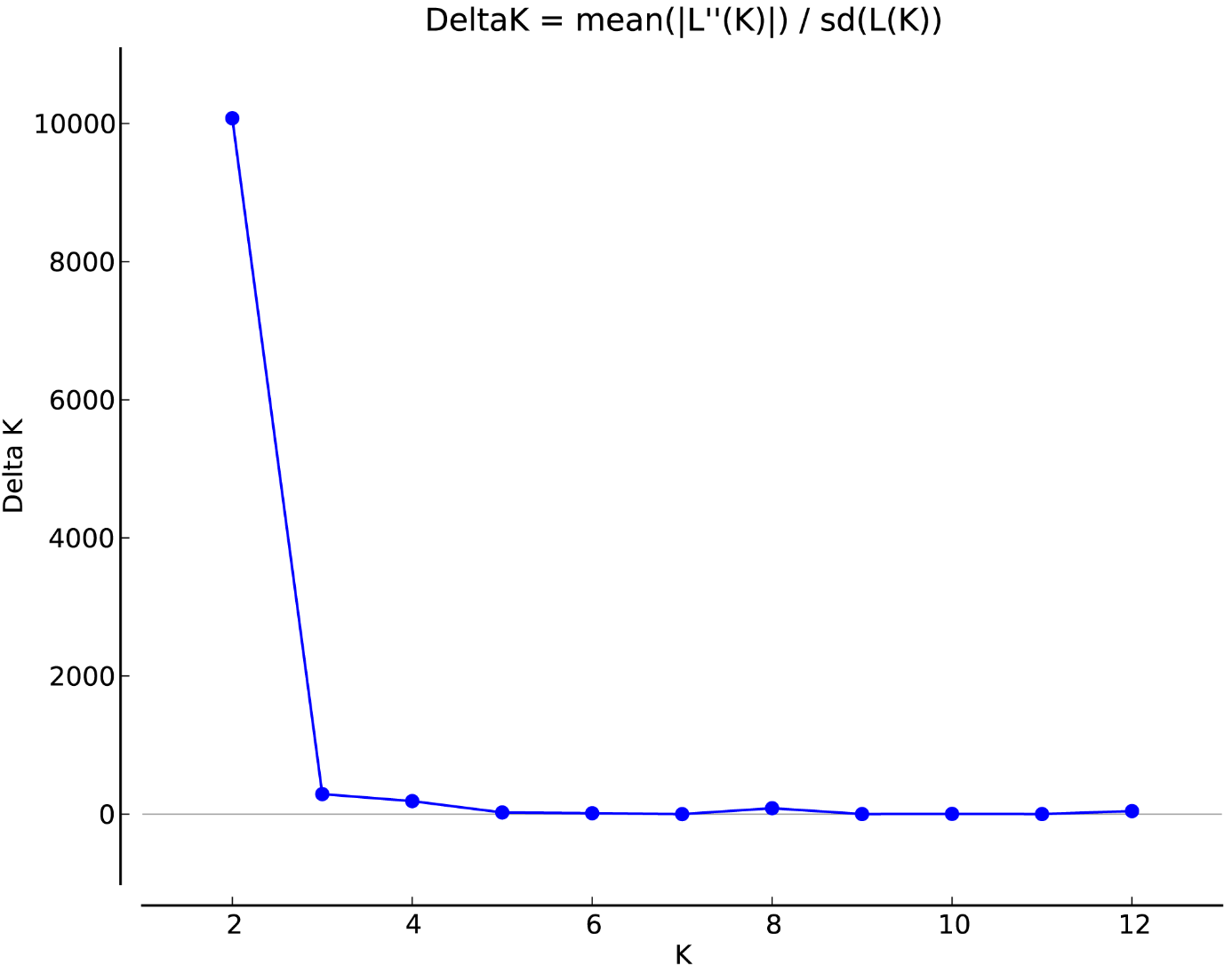

**Figure S2**. Plot of Delta K values for number of assumed subpopulations (K) ranging from 2 to 14.

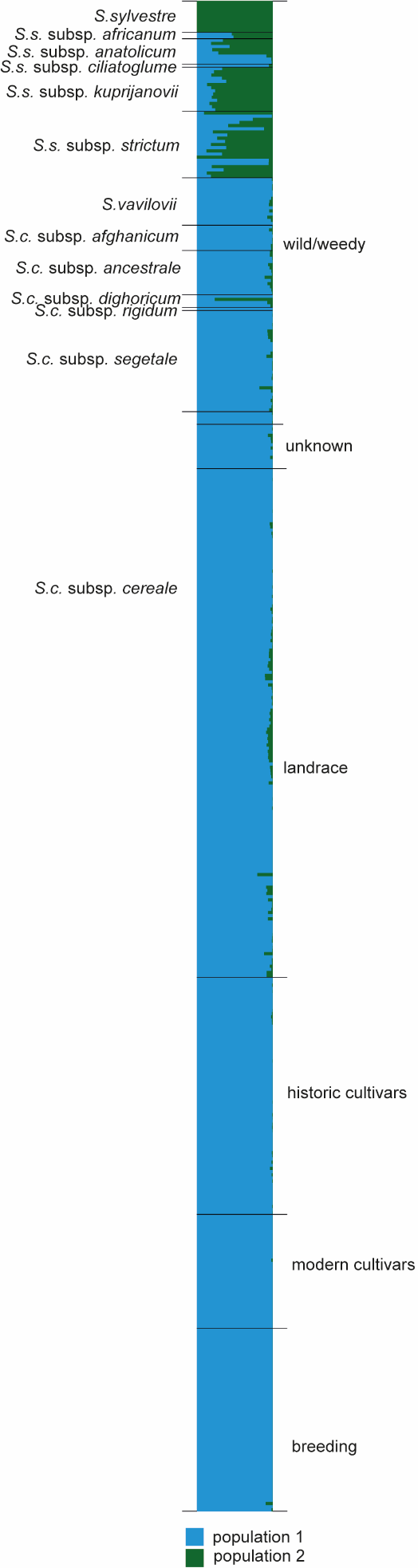

**Figure S3**. Population structure of 478 rye accessions at K=2 based on 12 846 SNPs. Each accessions is represented by a vertical stripe partitioned into coloured segments with lengths representing the membership fractions in the inferred clusters. The order of accessions in the plot is the same as in the Table S1.

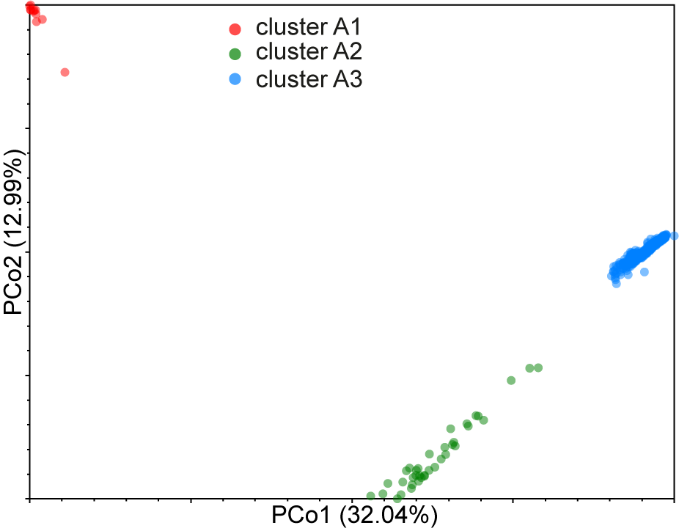

**Figure S4**. Principal Coordinates Analysis plot showing relationships between 478 rye accessions genotyped with 12846 SNPs with accessions labelled according to the outcome of NJ clustering.

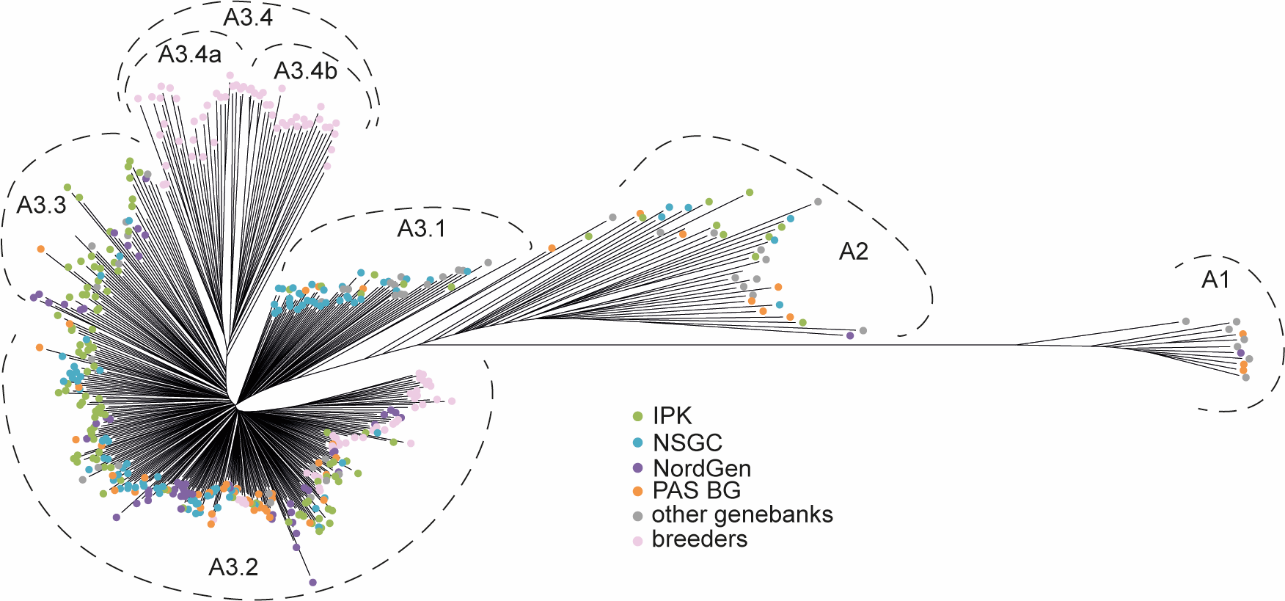

**Figure S5**. Neighbor-joining tree based on 12 846 SNP markers showing relationships between 478 rye accessions with accession labelled according to source.

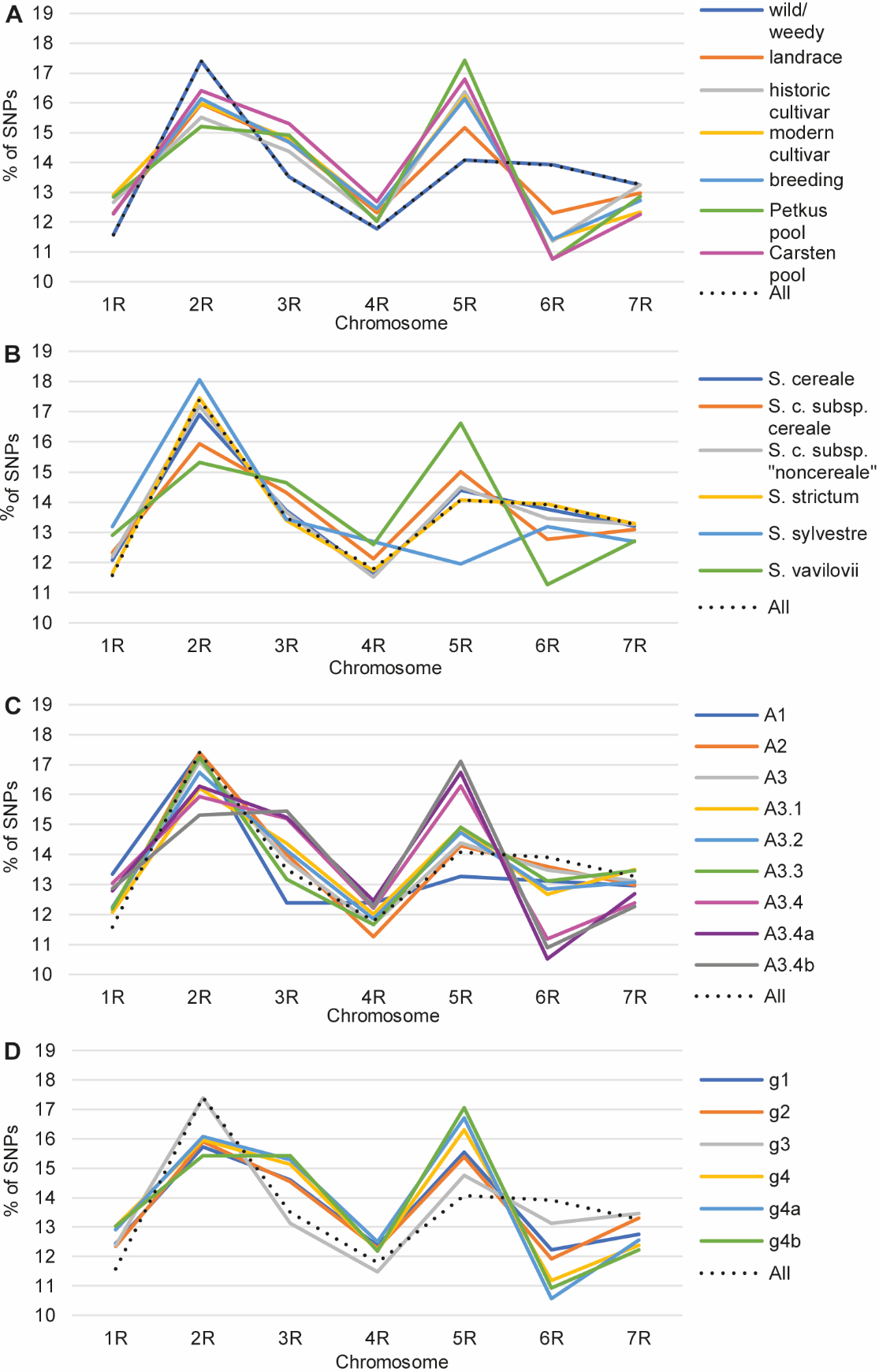

**Figure S6.** Chromosomal distribution of polymorphic SNPs by germplasm group. **A.** Accessions grouped according to improvement status. **B.** Accessions grouped according to taxonomy. **C.** Accessions grouped according to the outcome of NJ clustering. **D**. Accessions grouped according to their membership in sweep detection sets.

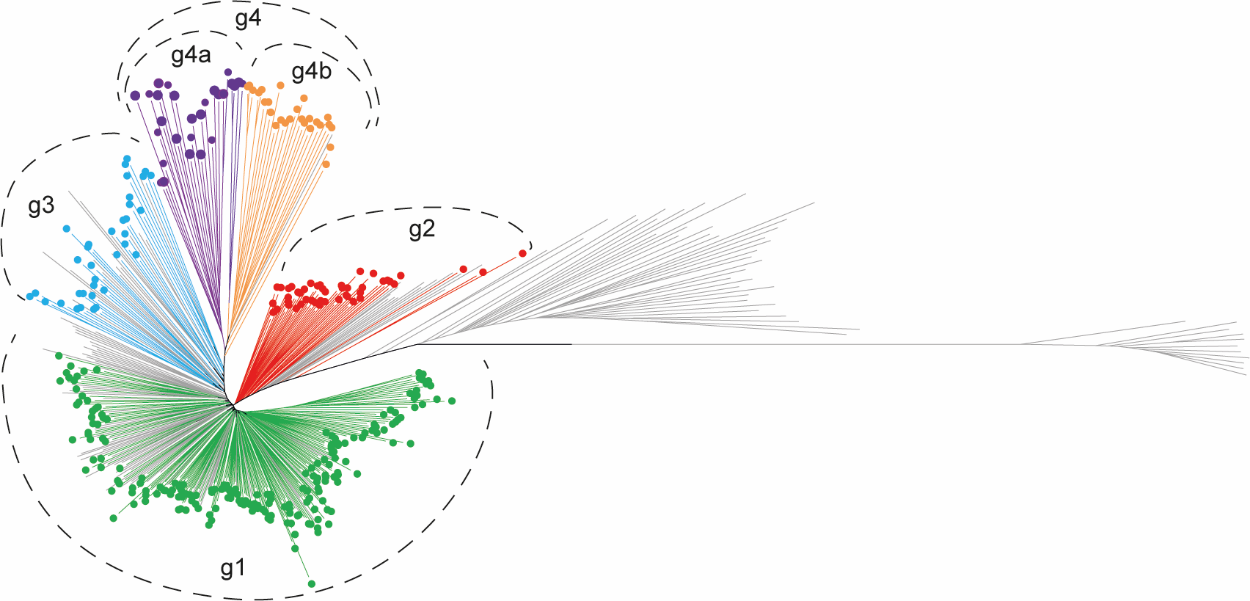

**Figure S7.** Neighbor-joining tree based on 12 846 SNP markers showing relationships between 478 rye accessions with branch colour indicating cultivated rye accessions belonging to the respective sweep detection set.
